# The essential molecular components for cellular CO_2_ sensing via connexins

**DOI:** 10.64898/2026.06.17.732653

**Authors:** Julien Pelletier, Jack Butler, Ariana Hassan, Nicholas Dale

**Author notes:** Contributed equally (Julien Pelletier, Jack Butler).

## Abstract

CO_2_ opens a subset of connexin hemichannels by binding to a site in the cytoplasmic domain of the channel. From outside the cell, CO_2_ must cross at least one membrane to reach this site. We have used Neuro-2A cells, which exhibit very low expression of CO_2_ permeable aquaporins (AQPs) and do not express any of the connexins (Cxs) known to be CO_2_ sensitive, to evaluate the minimal complement of molecular components required to recapitulate whole cell CO_2_ sensitivity mediated by connexins (assayed by either whole cell patch clamp recordings or real time recordings of ATP release via a co-expressed genetically encoded ATP sensor). Neuro-2A cells that expressed either Cx26, Cx32 or Cx43 on their own did not exhibit CO_2_-dependent connexin hemichannel gating. Expression of AQP1 or AQP5 either with or without carbonic anhydrase 2 (CA2) did not reveal any endogenous CO_2_ sensitivity of Neuro-2A cells. Only by expressing one of Cx26, Cx32 or Cx43 with either AQP1 or AQP5, plus CA2 were we able to reconstitute whole cell CO_2_ sensitivity. We found that expression of Cx26 with either AQP1 or AQP5 resulted in high levels of cell death. This was prevented by co-expression of CA2. Simulations of the influx and diffusion of CO_2_ show that CA2 prevents accumulation of intracellular CO_2_ and excessive activation of Cx26, thus protecting the cells from death. Surveying the transcriptome of cells that express CO_2_ sensitive connexins shows that many also express CO_2_ permeable aquaporins and CA2. We suggest that connexins, aquaporins and carbonic anhydrases represent the minimal trifecta of components required for cellular CO_2_ sensing.

## Introduction

CO_2_ opens hemichannels of five of the 21 connexins encoded in the human genome (Huckstepp et al., 2010; Meigh et al., 2013; Dospinescu et al., 2019; Dale, 2021; Butler and Dale, 2023; Dospinescu et al., 2025; Lovatt et al., 2025). CO_2_ binds to a conserved motif, the carbamylation motif, which is situated in the cytoplasmic loop (Meigh et al., 2013; Brotherton et al., 2022; Brotherton et al., 2024; Nijjar et al., 2025). CO_2_ forms a labile carbamate bond with the side chain primary amines of Lys residues within this loop (Nijjar et al., 2025). The carbamylated Lys residues then interact with positively charged residues (Lys or Arg) in the cytoplasmic domains of TM2 and TM4 of the neighbouring subunit (Meigh et al., 2013). Thus CO_2_, produced either systemically or from metabolically active cells, must cross at least one plasma membrane to reach the intracellular binding site on the connexin. Biological membranes, especially those that contain cholesterol, are poorly permeable to CO_2_ (Boron et al., 2011). This implies the need for the presence of channels that are permeable to CO_2_ such as a subset of the aquaporins (AQP0,1, 4, 5, 6 and 9) (Endeward et al., 2006; Boron, 2010; Geyer et al., 2013; Michenkova et al., 2021). Cells that sense and respond to CO_2_ might therefore be expected to co-express one of these aquaporins alongside a CO_2_ sensitive connexin. However, as aquaporins are always open, expression of CO_2_ permeable aquaporins would likely result in a steady state CO_2_ influx and accumulation of a cytosolic PCO_2_ sufficiently high to open any CO_2_ sensitive connexins that are also present. CO_2_ sensing cells must therefore be adapted to tolerate this high steady state CO_2_ influx. One possible adaptation is increased expression of carbonic anhydrase 2 (CA2), as this highly efficient cytosolic enzyme could counteract excessive intracellular accumulation of CO_2_ by converting it to carbonic acid which will spontaneously dissociate to bicarbonate and hydrogen ions.

To test these ideas, we have taken advantage of Neuro-2A cells which have very low expression of CO_2_ permeable aquaporins, and no expression of connexins known to be sensitive to CO_2_ (Kurosaki et al., 2021). Interestingly these cells do not express CA2 either (Kurosaki et al., 2021) and are thus an ideal model to investigate the need for combinatorial expression of connexins, aquaporins and CA2 to recapitulate whole cell CO_2_ sensitivity. We find that a combination of a CO_2_ sensitive connexin, a CO_2_ permeable aquaporin and CA2 must be expressed to reconstitute CO_2_ sensing. The presence of enhanced CA2 expression is required to protect the cell from the steady state CO_2_ influx, by ensuring that the steady state PCO_2_ within the cell remains sufficiently low to avoid significant opening of the connexin hemichannel under basal conditions.

## Results

### Recapitulation of connexin-dependent CO_2_ induced changes in whole cell conductance of Neuro-2A cells

Neuro-2A cells, owing to their very low expression of endogenous connexins are a favoured expression system for investigation of the properties of connexin hemichannels. Neuro-2A cells that had been treated with PEI transfection reagent without any cDNA (PEI control) did not show any whole cell conductance changes in response to CO_2_ (Fig 1). Expression of a CO_2_ sensitive connexin, Cx26, also failed to give any whole cell conductance changes. As Neuro-2A cells do not significantly express CO_2_ permeable aquaporins, we tested whether transfection of AQP1 might reveal endogenous CO_2_-dependent whole cell conductance changes. AQP1-transfected cells remained insensitive to CO_2_ (Fig 1). Co-transfection of AQP1 and Cx26 resulted in very substantial cell death (see later) so was not pursued as a viable strategy. We next co-transfected cells with a combination of AQP1 and CA2. Once again, these cells remained insensitive to CO_2_ (Fig 1). When a triple transfection of Cx26, AQP1 and CA2 was performed, cells reliably exhibited CO_2_ dependent whole cell conductance changes (Fig 1, Supplementary Fig 1). This was also true with the triple transfection of Cx43, another CO_2_-sensitive connexin, AQP1 and CA2. The whole cell conductance changes and resulting outward current were very similar to those previously reported for expression of either Cx26 or Cx43 in HeLa DH cells (Dospinescu et al., 2025; Nijjar et al., 2025), which happen to express CO_2_-permeable aquaporins and tolerate the expression of CO_2_-sensitive connexins.

**Figure 1:**
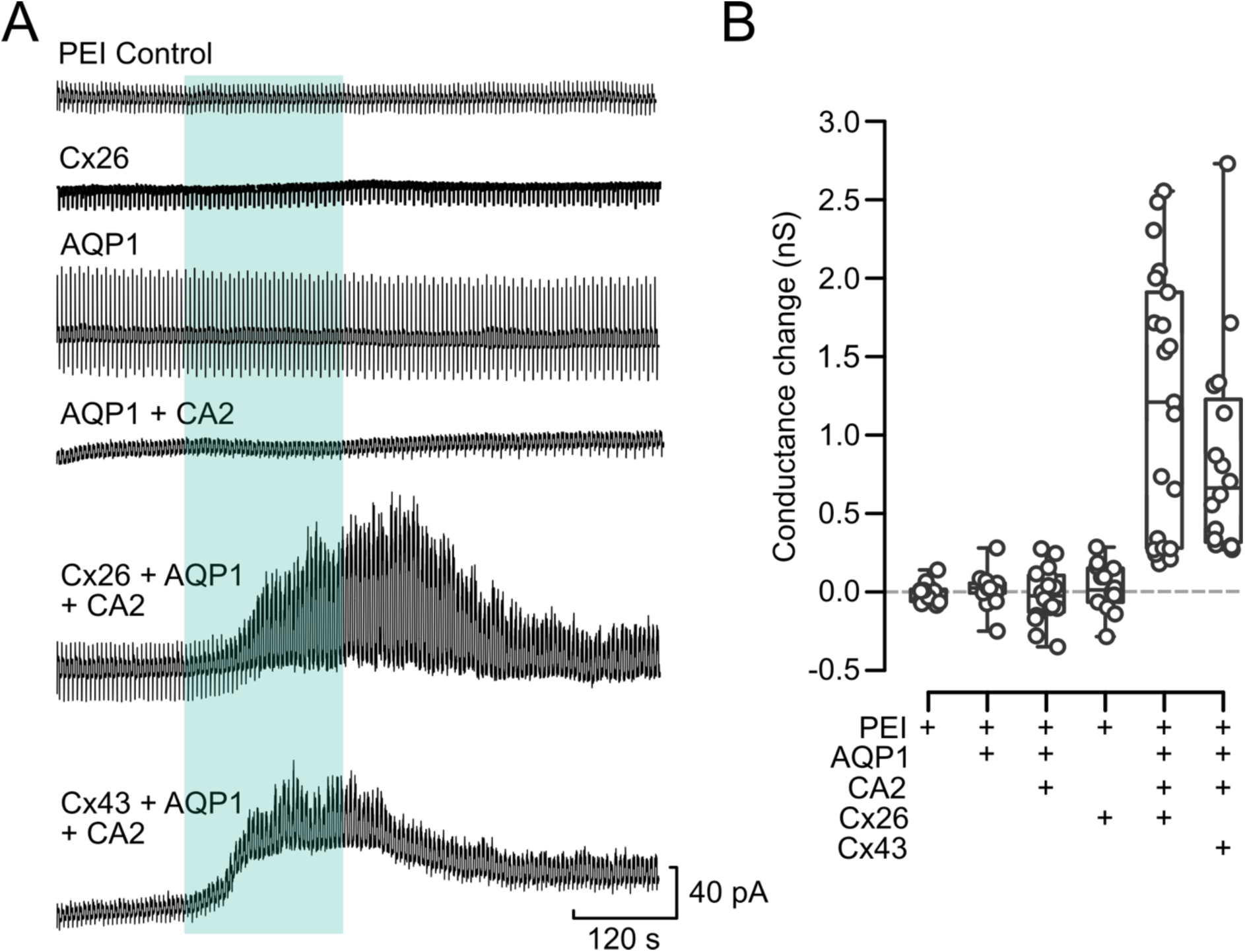
CO_2_ permeable aquaporins are required for CO_2_ sensing via connexins. **A)** Whole cell voltage clamp recordings from Neuro-2A cells under various transfection conditions. The cells were held at −50 mV and stepped to −30 mV every 5 s. The cells were bathed in aCSF with a PCO_2_ of 35 mmHg which was changed to 70 mmHg or 55 mmHg (Cx43 only) at the time indicated by the cyan rectangle. Treatment with the transfection reagent (PEI control, 2 independent treatments) or transfection with Cx26 (4 independent transfections), or AQP1 (2 independent transfections) or AQP1 + CA2 (2 independent transfections) did not cause any sensitivity to CO_2_. Only co-transfection of Cx26 or Cx43 plus AQP1 and CA2 (4 and 2 independent transfections respectively) gave robust CO_2_-triggered changes in whole cell conductance. **B)** Summary graph showing the data for all treatment groups. Each circle indicates a recording from one cell. The box represents the interquartile range, the bar the median, and the whiskers the range of the data. Kruskal-Wallis one way ANOVA: chi-squared = 63.245, df = 5, p = 2.59e-12.

### Recapitulation of CO_2_-dependent ATP release via connexins in Neuro-2A cells

By using the genetically encoded ATP sensor GRAB_ATP_, we have previously shown that hemichannels of Cx26, Cx32 and Cx43 are capable of mediating the release of ATP in response to small increases of PCO_2_ (Butler and Dale, 2023; Dospinescu et al., 2025; Lovatt et al., 2025; Nijjar et al., 2025). This assay is advantageous compared to patch clamp as the ATP release is unambiguously the result of a large-pore channel opening, and the fluorescence imaging assay can detect gating of CO_2_ dependent channels in many cells simultaneously. Expression of Cx26, Cx32 or Cx43 on their own (in combination with GRAB_ATP_) in Neuro-2A cells did not result in CO_2_-dependent ATP release (Fig 2, Supplementary Fig 2). We next expressed either AQP1 or AQP5 in combination with CA2 and GRAB_ATP_ in Neuro-2A cells. Once again, there was no evidence for CO_2_ evoked ATP release. The combination of Cx26 and CA2 did not result in CO_2_ evoked ATP release, although this combination often gave reductions in GRAB_ATP_ fluorescence (Fig 3, Supplementary Fig 3).

**Figure 2:**
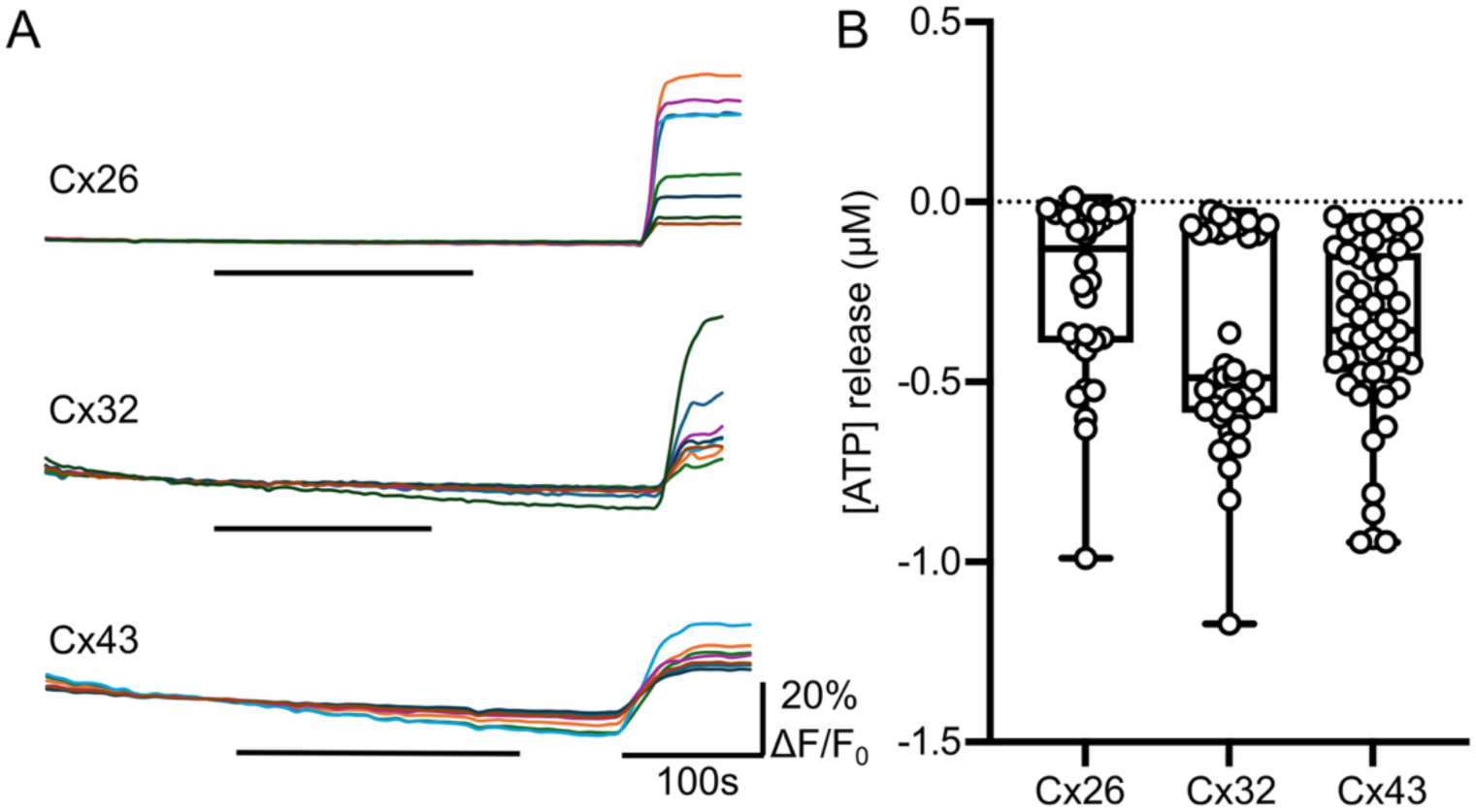
CO_2_ sensitive connexins expressed in Neuro-2A cells on their own do not permit CO_2_ dependent ATP release. **A)** Sample recordings showing GRAB_ATP_ fluorescence during the application of 55 mmHg PCO_2_ (Cx26 and Cx43) or 70 mmHg (Cx32) stimulus to cells expressing a CO_2_-sensitive connexin on its own. 3 µM ATP was applied at the end of the recording to calibrate the sensor. **B)** Box and whisker plots showing the lack of any CO_2_ evoked ATP release (5 independent transfections for each gene). The negative values are due to GRAB_ATP_ bleaching.

**Figure 3:**
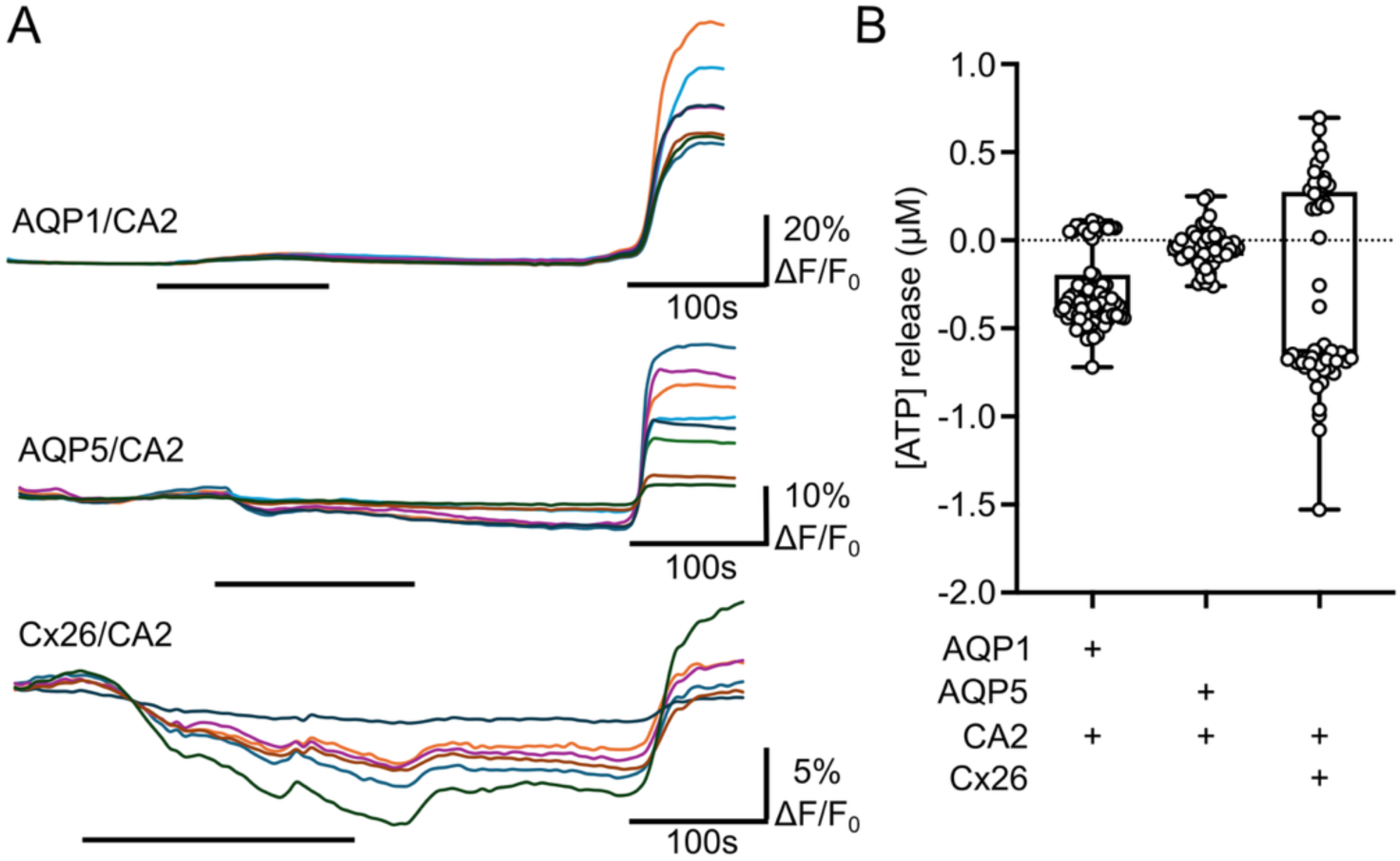
Co-expression of CO_2_ permeable aquaporins and carbonic anhydrase 2 (CA2) does not uncover CO_2_ sensitive responses in Neuro-2A cells. **A)** Sample recordings showing GRAB_ATP_ fluorescence during the application of 55 mmHg PCO_2_ stimulus to cells expressing either AQP1 and CA2, AQP5 and CA2 or Cx26 and CA2. 3 µM ATP was applied at the end of the recording to calibrate the sensor (5 independent transfections for each gene combination). **B)** Box and whisker plots showing the lack of any CO_2_ evoked ATP release. The negative values are due to GRAB_ATP_ bleaching.

Only the combination of a CO_2_ sensitive connexin (Cx26, Cx32 or Cx43), a CO_2_-permeable aquaporin (AQP1 or AQP5) and CA2 with GRAB_ATP_ recapitulated CO_2_-dependent ATP release from Neuro-2A cells (Fig 4, Supplementary Fig 4). The choice of CO_2_-permeable aquaporin made no difference to the amplitude of the CO_2_ evoked ATP release suggesting that they were interchangeable and both provided adequate rates of CO_2_ influx to support connexin hemichannel opening.

**Figure 4:**
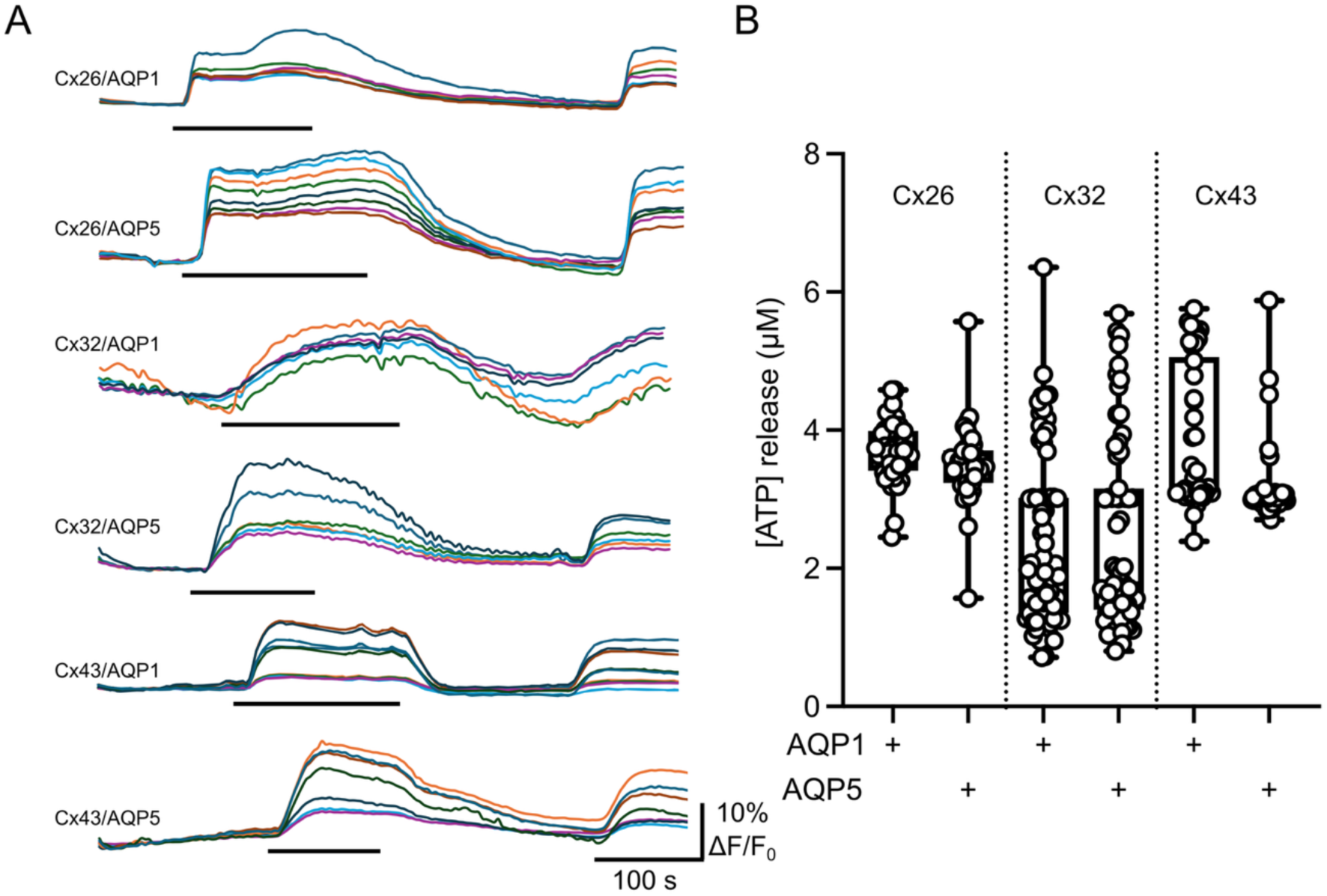
Combined expression of CO_2_ sensitive connexins plus CO_2_ permeable aquaporins and CA2 permit the reconstitution of CO_2_ sensitive ATP release. **A)** Co-expression of either AQP1 or AQP5 with one of Cx26, Cx32, Cx43 gave robust and reproducible ATP release in response to 55 mmHg (Cx26, Cx43 black bar) or 70 mmHg (Cx32, black bar). All cells co-expressed CA2. 3 µM ATP calibration applied at end of trace (5 independent transfections for each gene combination). **B)** Summary data showing that all combinations of AQP1 or AQP5 with one of the CO_2_-sensitive connexins (plus CA2) gave CO_2_-evoked ATP release.

### Expression of carbonic anhydrase is essential to avoid cell death

Our observations suggested that certain combinations of gene expression increased the death of the Neuro-2A cells. We therefore quantified the proportion of dead cells 48 h after transfection compared to the PEI controls. Expression of Cx26, AQP1 or CA2 on their own had very little effect on cell death compared to the controls (Fig 5A). However, the expression of Cx26 plus AQP1 gave an approximately 20-fold increase in the number of dead cells after 48 h, which was almost completely prevented by co-expression of CA2 (Fig 5A). We also examined the effect of the different combinations of gene expression on total (dead and living) cell numbers (i.e. growth from 24 to 48 h, Fig 5B). The expression of Cx26 or AQP1 did not prevent the increase in cell numbers from 24 to 48 h that was seen in the PEI control. Expression of CA2 on its own, however, prevented this increase in cell number from 24 to 48 h. In cells that expressed either Cx26 plus AQP1 or Cx26 plus AQP1 plus CA2 no increase in cell numbers was evident from 24 to 48 h post transfection. Thus, although the co-expression of CA2 alongside Cx26 and AQP1 prevented the massive increase in cell death caused by Cx26 and AQP1 co-expression, it did not restore the capacity of these cells to proliferate. Presumably there are yet further adaptations that enable cells to tolerate the co-expression of CO_2_ sensitive connexins and CO_2_ permeable aquaporins.

**Figure 5:**
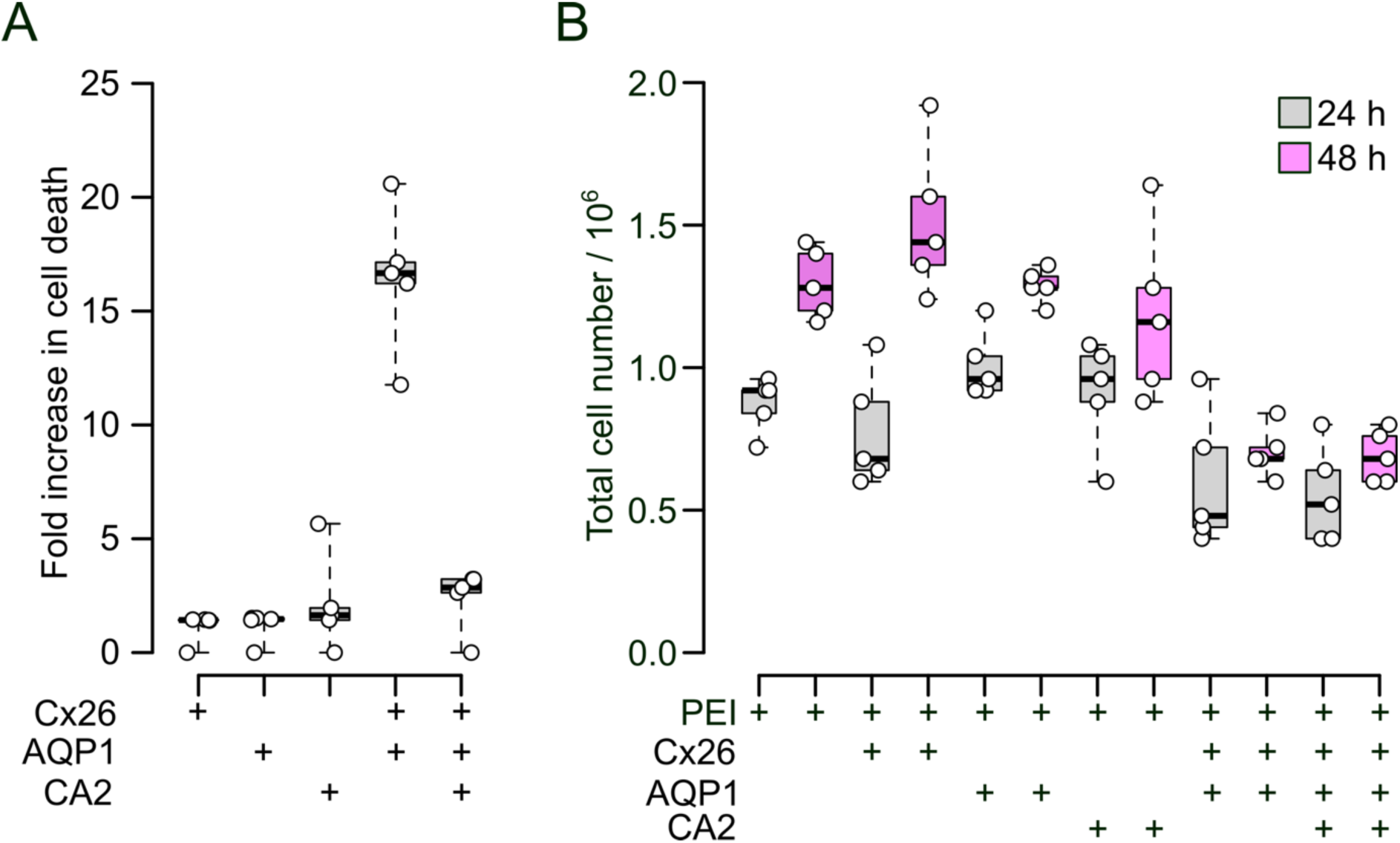
Co-expression of CO_2_ sensitive connexins and CO_2_ permeable aquaporins results in cell death that is prevented by co-expression of CA2. **A)** The effect of gene expression on the survival of Neuro-2A cells in a CO_2_-incubator (5% CO_2_, 37 °C) expressed as the fold-change in cell death compared to the PEI treated controls. Kruskal-Wallis Anova: chi-squared = 14.759, df = 4, p = 0.005. Mann-Whitney U test: Cx26+AQP1 vs Cx26+AQP1+CA2, p = 0.012. **B)** The change in total cell number (live and dead) over 24 h under different transfection conditions. The PEI control and transfections with Cx26 or AQP1 exhibit a substantial increase in cell number over the 24 h period. Transfection with CA2 alone or in combinations with Cx26 and AQP1 supress cell proliferation, Expression of Cx26 and AQP1 alone suppresses cell proliferation. Mann-Whitney U tests: PEI 24 vs 48, p = 0.012; Cx26 24 vs 48, p = 0.008; AQP1 24 vs 48, p = 0.015; CA2 24 vs 48, p = 0.207; Cx26+AQP1 24 vs 48, p = 0.462; Cx26+AQP1+CA2 24 vs 48, p = 0.246. N=5 independent transfections of cells, for A and B.

### Simulations show the importance of carbonic anhydrase in maintaining low cytosolic PCO_2_

To understand better the role of CA2, we simulated the diffusion of CO_2_ in a 20 x 20 µm plane with a continual influx of CO_2_ (PCO_2_ of 90 mmHg/ms, equivalent to 2.7 mM/ms) at its centre. Throughout the plane CA2 was present and modelled with Michaelis-Menten kinetics (K_m_ of 13 mM for CO_2_, from the literature). Cx26 hemichannels were also present throughout the plane and the CO_2_-dependent gating of these channels was simulated via the Hill equation with parameters drawn from the literature (EC_50_ of 40 mmHg, Hill coefficient of 4). When the V_max_ of CA2 was set to 1.5 mM/s, after 200 ms of simulation, PCO_2_ rapidly decayed from a maximum value of just under 70 mmHg at the point source, to a very low level of 1-2 mmHg outside of a radius of a few microns from the point source. Consequently, the activation of Cx26 hemichannels was restricted to a narrow circular area around the point source with a radius of ∼2 µm. When the V_max_ of CA2 was reduced to 1.5 µM/s, the PCO_2_ at the point source was attenuated less (peak of ∼75 mmHg) and decayed from the point source to a baseline of about 30 mmHg across the plane. This baseline PCO_2_ was sufficient to give substantial opening of Cx26 (∼25 % of maximal conductance) across the entire plane (Fig 6). This modelling suggests that CA2 has relatively little effect on the peak value of PCO_2_ at the point source but a very large effect on the steady state baseline PCO_2_ away from the point source. This explains why expression of a CO_2_ sensitive connexin and a CO_2_ permeable aquaporin without compensatory expression of CA2 results in cell death. The build-up of PCO_2_ and consequent steady state gating of the connexin will impose on the cell a very high metabolic load simply to restore transmembrane ionic gradients, let alone to replenish the continual efflux of key cytosolic metabolites such as ATP and lactate.

**Figure 6:**
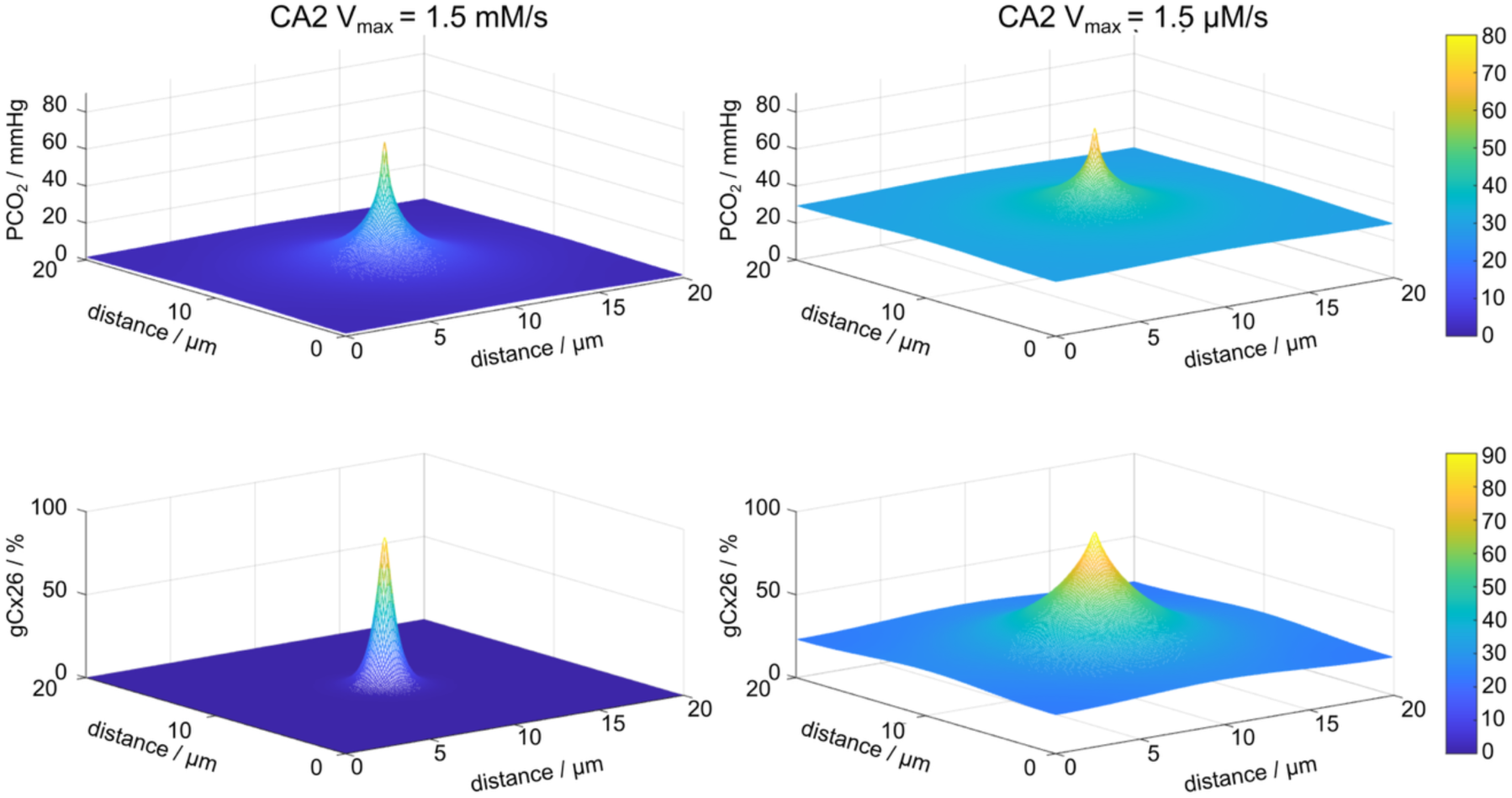
Simulations of CO_2_ entry and diffusion in a 2D plane show the importance of carbonic anhydrase. The simulation is based on the partial differential equation: ∂**CO_2_**/∂t = D∇^2^**CO_2_** + J_i_ - V_max_**CO_2_**/(**CO_2_**+K_m_) - K_f_**CO_2_**. Where D is the aqueous diffusion coefficient for CO_2_ (2.2 µm^2^/ms), ∇^2^ is the double differential with respect to distance in the x and y dimensions, **CO_2_** is a matrix for the concentration of CO_2_ (in x and y dimensions), J_i_ the influx of CO_2_ at the point source (90 mmHg/ms), V_max_ and K_m_ (set to 13 mM) are the parameters for the Michaelis-Menten equation describing CA2, and K_f_ is the rate constant for the spontaneous hydration of CO_2_. The gating of Cx26 hemichannels was described by the Hill equation with an EC_50_ of 40 mmHg and a Hill coefficient of 4. These equations were solved numerically using code written for Matlab.

### Co-expression of CO_2_-sensitive connexins, CO_2_-permeable aquaporins and carbonic anhydrases in non-neuronal brain cells

In the brain, CO_2_ sensitive connexins are expressed predominantly in non-neuronal cells. We therefore explored the human single brain cell transcriptome (Siletti et al., 2023), analyzing expression in pericytes, microglia and astrocytes (Fig 7). A relatively small proportion of pericytes coexpressed either Cx43 (GJA1) or Cx30 (GJB6) with AQP1 or AQP4. In pericytes there was a very high level of expression of CA2 meaning that virtually all cells that expressed Cx43 also expressed CA2 (Fig 7). For microglia there was a substantial proportion of cells that expressed Cx43 or Cx32 (GJB1) with either AQP1 or AQP4. A similarly substantial proportion of Cx43 or Cx32 expressing cells also expressed CA2 (Fig 7). For astrocytes, virtually all cells that expressed either Cx43 or Cx30 also expressed AQP4. A subset of astrocytes coexpressed AQP1 with Cx43 or Cx30. There was also widespread co-expression of CA2 in virtually all Cx43 expressing cells (Fig 7).

**Figure 7:**
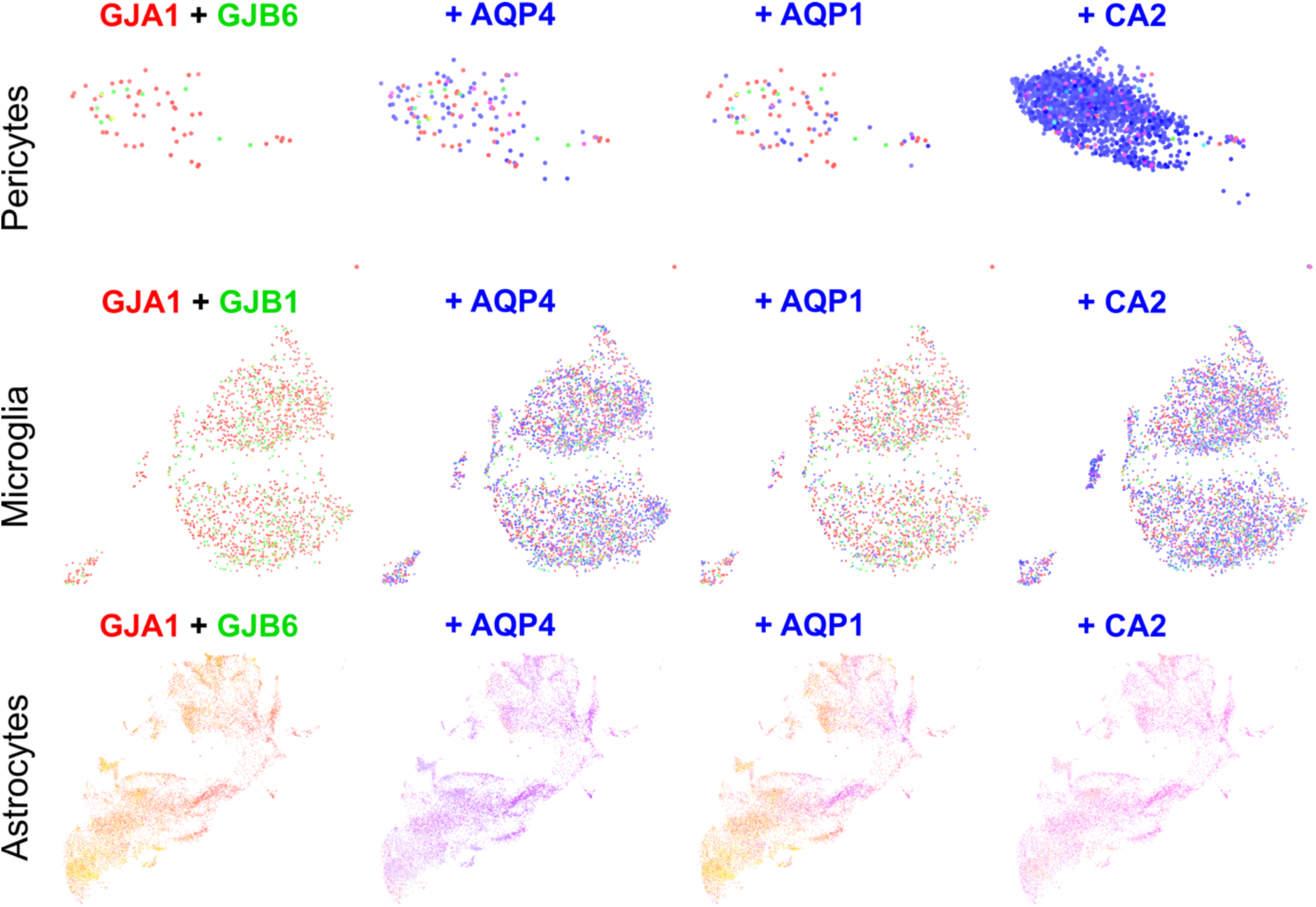
Expression of CO_2_ sensitive connexins, CO_2_ permeable aquaporins, and CA2 in human pericytes, microglia and astrocytes. Analysis via the Allen Brain Cell Atlas of scRNAseq data from Linnarsson, Lein, Bakken (2023) Transcriptomic Characterization of Cell Types in Human Brain. Available from https://assets.nemoarchive.org/dat-5ie1mec. For each cell type the predominant CO_2_ sensitive connexins were mapped in combinations with the predominant CO_2_ permeable aquaporins and CA2.

**Figure 8:**
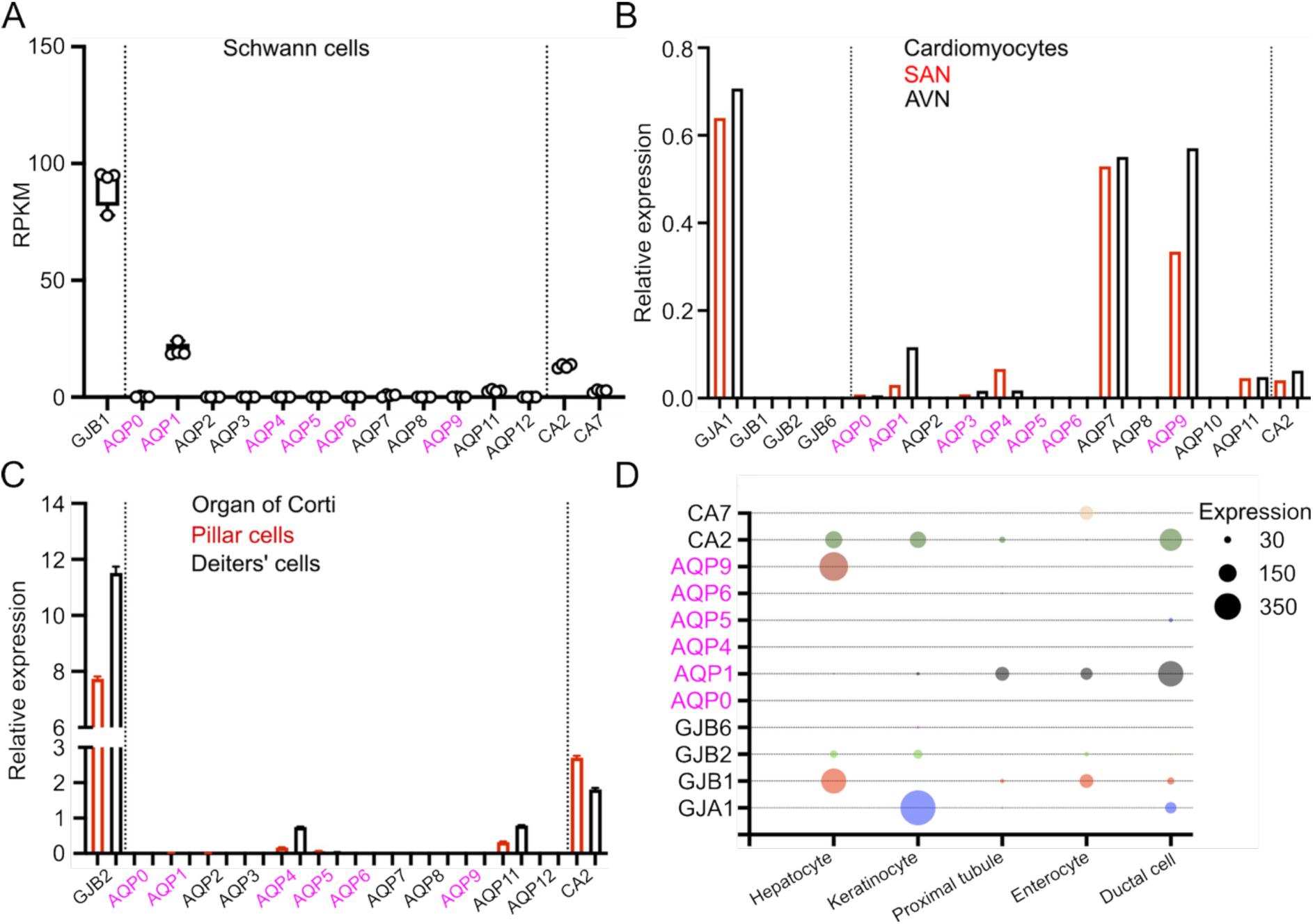
Cellular expression of the CO_2_ sensing trifecta in peripheral tissue. A) scRNAseq data for myelinating Schwann cells from the Sciatic Nerve Atlas (SNAT) (Gerber et al., 2021). Expression plotted as reads per kilobase of transcript per million (RPKM). B) scRNAseq data for cardiomyocytes from the sinoatrial node (SAN) and atrioventricular node (AVN) from the Heart Cell Atlas (Kanemaru et al., 2023). C) scRNAseq data for pillar cells and Deiters’ cells of the Organ of Corti from NCBI Gene Expression Omnibus GSE111347 (Liu et al., 2018). D) scRNAseq data for liver hepatocytes, suprabasal keratinocytes, kidney proximal tubule cells, small intestine enterocytes and pancreatic ductal cells from the Human Protein Atlas (https://www.proteinatlas.org) (Uhlén et al., 2015). Expression levels are in normalized counts per million (nCPM).

### Co-expression of CO_2_-sensitive connexins, CO_2_-permeable aquaporins and carbonic anhydrases in peripheral tissues

To extend our analysis beyond the brain, we examined single cell transcriptomic data from Schwann Cells of peripheral myelin, support cells of the organ of corti, suprabasal keratinocytes of skin, proximal tubule cells of kidney, liver hepatocytes, ductal cells of pancreas, enterocytes of small intestine and cardiac myocytes.

In peripheral myelin we have already shown an important role for AQP1 and CA2 in the CO_2_ mediated gating of Cx32 in Schwann Cells. The Schwann Cell transcriptome shows that these three components are expressed and indicates that the expression level of AQP1 and CA2 is considerably less than that of Cx32 (approximate ratio 4:1:1).

In the support cells of the organ of corti Cx26 was strongly expressed in the Pillar and Dieter’s cells, but only the Dieter’s cells had the complete CO_2_ sensing trifecta with significant AQP4 expression. Cardiomyocytes from the atrioventricular node (AVN) and sinoatrial modes (SAN) expressed the complete trifecta (Cx43, AQP9 and CA2). Pancreatic ductal cells, kidney proximal tubule cells, enterocytes from the small intestine all expressed the trifecta with either AQP1 or AQP9 being the main CO_2_ conduit. An exception seems to be suprabasal keratinocytes which have very high levels of Cx43 and Cx26 expression but negligible expression of CO_2_ permeable aquaporins.

## Discussion

### Neuro-2A transcriptome

As our study relied on the serendipitous discovery that connexin expression alone in Neuro-2A cells is not sufficient to endow them with whole cell CO_2_ sensitivity, it is instructive to examine the transcriptome of Neuro-2A cells in the dataset (NCBI Gene Expression Omnibus GSE180136) created by Kurosaki et al. (2021) to assess the expression of relevant genes (aquaporins, carbonic anhydrases and various transporters relevant to the maintenance of intracellular pH, summarised in Table 1). The aquaporins with strongest expression (AQP3 and AQP11) are not permeable to CO_2_. Although there is weak expression of AQP1 and AQP5 this is evidently not enough to allow activation of Cx26, Cx32 or Cx43 hemichannels by changes of extracellular PCO_2_. Interestingly CA2 is not expressed by the Neuro-2A cells, but they do express CA7 which has a cytosolic localisation and has high levels of catalytic activity. Nevertheless, our results imply that this level of expression of CA7 is not sufficient to protect the Neuro-2A cells when CO_2_ sensitive connexins and CO_2_ permeable aquaporins are co-expressed on their own without additional expression of CA2. CA11, also expressed by Neuro-2A cells, is not catalytically active so its function remains unclear. CA11 is strongly expressed in some brain regions therefore its expression in the Neuro-2A cells may not be unexpected. One of the consequences of higher CO_2_ influxes and carbonic anhydrase activity will be the creation of hydrogen and bicarbonate ions. In this regard it is interesting that Neuro-2A cells have very strong expression of the Na^+^-H^+^ exchanger (Slc9a1). They also have very strong expression of Slc4a2, an anion exchanger that could swap Cl^-^ and HCO_3_^-^ ions. However, expression of Slc4a10, the Na^+^ linked HCO_3_^-^ transporter is only weak. On this basis we expect Neuro-2A cells could regulate their pH in the face of overexpression of a CO_2_ permeable aquaporin and CA2. However, our data suggests that this does compromise growth and it is possible that enhanced expression of some other component is required such as Slc4a10.

**Table 1:** Expression of genes relevant to CO_2_ sensitivity and pH regulation in Neuro-2A cells. Based on data from NCBI Gene Expression Omnibus GSE180136. The classification of strength of expression level is based on the raw read counts (RRC) and is: Very weak, RRC≤5; Weak, 5<RRC≤25; Medium, 25<RRC≤100; Strong,100<RRC≤1000; and Very strong, RRC>1000.

| Category | Gene name | Expression level |
| --- | --- | --- |
| **Aquaporins** | AQP0, AQP4, AQP6, AQP9 | Very weak |
|  | AQP1, AQP5 | Weak |
|  | AQP11 | Medium |
|  | AQP3 | Strong |
| **Carbonic anhydrases** | Ca2 | Very weak |
|  | Ca6, Ca7, Ca11 | Strong |
| **Na<sup>+</sup>/H<sup>+</sup> exchange** | Slc9a1 | Very strong |
| **Anion exchangers** | Slc4a1, Slc4a4 | Very weak |
|  | Slc4a2 | Very strong |
| **Na<sup>+</sup>-linked HCO<sub>3</sub><sup>-</sup> transport** | Slc4a10 | Weak |

### Required components for whole cell CO_2_ sensitivity

Our results show that co-expression of a CO_2_ permeable aquaporin and CA2 alongside a CO_2_ sensitive connexin is required to recapitulate CO_2_ sensing in the Neuro-2A cells. The aquaporin is required to allow the CO_2_ to reach its binding site on the cytoplasmic loop of the connexin. These data support the growing body of literature that suggests that certain aquaporins are necessary to allow significant permeation of CO_2_ across biological membranes. Arguably, the aquaporins and the connexins could be regarded as a CO_2_ sensing duopoly, with both being essential even though the critical transducing step (carbamate bond formation) occurs on the connexin. The role of CA2 is to lower the resting PCO_2_ to prevent continual activation of the connexin hemichannel and thus an unsustainable metabolic load that leads to the death of the cell. The actions of CA2 also maintain a steep transmembrane diffusion gradient for CO_2_ influx via the aquaporins further facilitating CO_2_ sensitivity via connexins. There is experimental support for CA2 enhancing CO_2_ fluxes through aquaporins (Wang et al., 2026). Thus, CA2 does not have a direct role in CO_2_ detection but sets the conditions that make it possible. It is also possible that other cytosolic isoforms of CA, notably CA1, CA3, CA7 and CA13, could perform a similar role to CA2 in facilitating whole cell CO_2_ sensitivity.

Modelling the diffusion of CO_2_ and the effect of CA2 shows that this enzyme (if uniformly present in cytosol) effectively limits the activation of CO_2_ sensitive connexins to a radius of less than 3 µm from a point source of CO_2_ entry. This in turn implies that CO_2_ sensitive connexins and CO_2_ permeable aquaporins must be closely co-located in the membrane to permit effective whole cell CO_2_ sensing. We have recently obtained evidence for this proximity in the paranodes of Schwann Cells that form peripheral myelin (Butler et al., 2025) and in the terminals of Dorsal raphe neurons that synapse onto dopaminergic neurons in the ventral tegmental area (Huang et al., 2025).

### Co-expression of the essential molecular components of CO_2_ sensing in brain

Our data show that cellular expression of a CO_2_ sensitive connexin is not sufficient on its own to endow cellular CO_2_ sensitivity: there must also be expression of a CO_2_ permeable aquaporin and a carbonic anhydrase to complete the CO_2_ sensing trifecta.

Analysis of the human brain cell transcriptome and focussing on non-neuronal cells, shows that virtually all astrocytes possess the CO_2_ sensing trifecta. This implies that almost all astrocytes have the capacity to sense and react to changes in PCO_2_. We have demonstrated this for astrocytes in the hippocampus (Dospinescu et al., 2025). A substantial proportion of microglia also express the CO_2_ sensing trifecta, again implying that they are CO_2_ sensitive. Conceivably, sensitivity to CO_2_ may assist migration of microglia to metabolic hotspots that could be useful for their roles as immune cells in the brain. In pericytes there was relatively little expression of the complete CO_2_ sensing trifecta, suggesting that most pericytes would be insensitive to changes in PCO_2_. This is perhaps surprising given their role in controlling cerebral blood flow (Peppiatt et al., 2006), where local changes in PCO_2_ might be a relevant physiological variable (Hosford et al., 2022).

There are further examples of the complete CO_2_ sensing trifecta in neural cells where connexin-mediated CO_2_ sensitivity has important physiological roles. In peripheral myelinated nerves both the axonal membrane and Schwann cell membrane express AQP1 (Ulrich et al., 2014; Segura-Anaya et al., 2015; Butler et al., 2025), and this allows CO_2_ produced in the axon to activate Cx32 hemichannels in the Schwann cell (Butler et al., 2025). Our analysis of the Schwann Cell transcriptome supports these observations. The terminals of Dorsal Raphe serotonergic neurons that innervate the dopaminergic neurons of the Ventral Tegmental Area co-express Cx26 and AQP5 permitting CO_2_ dependent 5-HT release from these terminals thus contributing to the hypercapnic arousal reflex (Huang et al., 2025).

### Co-expression of the essential molecular components of CO_2_ sensing in peripheral tissues

Probing scRNA-seq data from different tissues suggests that the CO_2_ sensing trifecta is widespread. Our analysis predicts cellular CO_2_ sensitivity in cardiomyocytes, pancreatic ductal cells, enterocytes of the small intestine, hepatocytes and kidney proximal tubule cells. In the organ of corti we predict CO_2_ sensitivity of the Deiters’ cells but not the pillar cells. Deiters’ cells are in close association with the outer hair cells which express prestin which enables their electromotility. This in turn amplifies the vibrations of the basilar membrane and enhances the sensitivity of hearing at low sound pressure levels (Dallos et al., 2008). Hyperpolarisation of Dieters’ cells alters the electromechanical properties of the basilar membrane (Lukashkina et al., 2024). If Deiters’ cells really are CO_2_ sensitive, then ambient CO_2_ in the organ of corti could alter the sensitivity of hearing.

Interestingly, suprabasal keratinocytes do not appear to express the complete trifecta suggesting that cellular CO_2_ sensitivity in skin may be unlikely. Additionally, Cx50 which is CO_2_ sensitive, and AQP0 (also known as Mip) are coexpressed in lens fibre cells (Berthoud and Ngezahayo, 2017; Li et al., 2022), which display CO_2_ sensitivity. The coordinated expression of CO_2_ permeable aquaporins and CO_2_ sensitive connexins plus CA2 is thus a widespread phenomenon in both brain and peripheral organs and supports the hypothesis that the modulatory/signalling actions of CO_2_ mediated by connexins are a universal feature of vertebrate physiology.

### Implications for the study of CO_2_ permeable channels

The CO_2_ permeability of aquaporins is a relatively recent discovery and is thought to occur through a combination of the pores within each monomer and the pore that is formed at the interface of the aquaporin subunits in the centre of the homotetramer (Michenkova et al., 2021; Musa-Aziz et al., 2026; Shinn et al., 2026). Assays of aquaporin CO_2_ permeability have relied on detecting the CO_2_ flux via a change in pH at the membrane surface (Musa-Aziz et al., 2009). We suggest that using Neuro-2A cells, expressing a CO_2_ sensitive connexin such as Cx26 with GRAB_ATP_ as an assay of hemichannel gating may be a technically advantageous system for examining the mechanisms of CO_2_ permeability of aquaporins and any other potential transmembrane conduits for CO_2_ such as the Rhesus factor (Endeward et al., 2008).

## Methods

### Plasmids and constructs used

pDisplay-GRAB_ATP1.0-IRES-mCherry-CAAX was a gift from Yulong Li (Addgene plasmid # 167582; http://n2t.net/addgene:167582; RRID:Addgene_167582).

pCMV-BI was a gift from Bill Hansson (Addgene plasmid # 126475; http://n2t.net/addgene:126475; RRID:Addgene_126475).

Initially we used single genes per plasmid. But as the number of genes increased, we used a bicistronic plasmid, pCMV-BI, that permits expression of two genes, one under a maain promoter and the other under a minimal promoter. In the patch clamp recordings we combined Cx26 or Cx43 (main promoter) with AQP1/AQP5 (minimal promoter), and then separately expressed CA2-eGFP from the pcDNA plasmid. For the GRAB_ATP_ recordings, for convenience and flexibility, we finally settled on putting CA2 and the AQPs respectively under the main and minimal promoters of pCMV-BI and then transfecting the relevant connexins (Cx26, Cx32 or Cx43) with a pCAG based construct.

### Cell culture

Neuro2a cells (RRID:CVCL_0470) were grown in low-glucose Dulbecco’s modified eagle medium (DMEM) (D6046) supplemented with 10% FBS and 50μg/mL penicillin/streptomycin. The Neuro2a cells were plated onto coverslips at a density of 7.5 x 10^4^ cells per well of a 6 well plate

### Transfection

For patch clamp recordings cells to express single genes 1 µg of the relevant construct in 3 µl of PEI was used to transfect the cells. To express multiple genes, cells were transiently transfected using a mixture of 500 ng of the pCMV-BI-Cx-AQP construct and 500 ng of the pcDNA-CA2 construct and 3 μl PEI for 6-8 hrs. recordings were taken 24-48 hours after transfection.

For GRAB_ATP_ recordings, to evaluate the complete CO_2_ sensing transfecta, cells were transiently transfected using a mixture of 250 ng of the pCMV-BI-CA2-AQP construct, 250 ng of the pCag-Cx-mCherry construct and 500 ng of the GRAB_ATP_ and 3 μl PEI for 6-8 hrs. Evaluation of individual genes or pairs of genes was performed with individual expression plasmids from Table 2. The total amount of DNA transfected was always 1 µg in 3 µl of PEI. Cells were imaged 48 hours after transfection. We used a protocol to measure ATP release from cells developed and described in our previous work (Butler and Dale, 2023).

**Table 2:** Constructs used to express connexin. Aquaporin and CA2 genes in Neuro-2A cells.

| Gene | Plasmid | Uniprot accession | Tag |
| --- | --- | --- | --- |
| Cx26 | pCAG | P29033 | mCherry |
| Cx32 | pCAG | P08034 | mCherry |
| Cx43 | pCAG | P17302 | mCherry |
| CA2 | pcDNA3.1 | P00918 | eGFP |
| AQP1 | pCAG | P29972 | mRuby |
| AQP5 | pCAG | P55064 | mRuby |
| Cx26/Cx43<br>(main promoter) | pCMV-BI |  | mTurquoise2 |
| AQP1/AQP5<br>(minimal promoter) |  |  | mRuby |
| CA2 (main promoter) | pCMV-BI |  | No tag |
| AQP1/AQP5<br>(minimal promoter) |  |  | mRuby |

### Fluorescence imaging

48 hours after transfection, cells were perfused with control aCSF (35 mmHg) until a stable baseline was reached, before switching to either hypercapnic aCSF. Once a stable baseline was reached after solution change, cells were returned to perfusion with control aCSF (35 mmHg). Recordings were calibrated by application of 3 μM ATP.

All cells were imaged by epifluorescence (Scientifica Slice Scope, Cairn Research OptoLED illumination, 60x water Olympus immersion objective, NA 1.0, with either a Hamamatsu ImagEM EM-CCD camera, Metafluor software, or Tucsen Libra 22 sCMOS camera controlled by Micromanager). GRAB_ATP_ was excited by a 470 nm LED, with emission captured between 504-543 nm. C-terminal mCherry or mRuby tags, which were excited by a 535 nm LED and emission captured between 570-640 nm. Only cells expressing both GRAB_ATP_ and mCherry were selected for recording, with GRAB_ATP_ images acquired every 4 seconds.

### Analysis of GRAB_ATP_ fluorescence

Analysis of GRAB_ATP_ was performed in ImageJ (Schneider et al., 2012). Cell recordings were corrected for any motion using the Image Stabilizer plugin (Li, 2008). For cells that expressed all of the required constructs, an ROI was drawn around the GRAB_ATP_ membrane expression and median fluorescence measured for each image. The fluorescence pixel intensity (F) was normalised to the baseline fluorescence (F_0_). The change in normalised fluorescence (ΔF/F_0_) evoked via hypercapnia was recorded for each cell.

We converted changes in normalised fluorescence evoked by hypercapnia into the concentration of ATP released by normalising them to the ΔF/F_0_ produced by a 3 μM ATP calibration solution. Over this range, the calibration curve for GRAB_ATP_ is approximately linear (Wu et al., 2022; Butler and Dale, 2023). Statistical comparisons were performed considering each cell as an independent replicate. Five transfections were performed.

### Cell viability assay

Cells plated at a density of 1 × 10^5^ cells per well in a six-well plate were transfected as described. After 48 h, the media was collected, and a PBS wash also collected in the same tube. Cells were then harvested with trypsin and added to the tube containing the media and PBS. The cells were then pelleted and resuspended in 2 ml of media. An aliquot was mixed 1 : 1 with trypan blue. Using a haemocytometer, the cells were counted to determine the proportion of dead (blue) cells. This was expressed relative to the proportion of dead cells found in similar cultures of dummy-transfected Neuro-2A cells, that were treated with PEI but no DNA. Five transfections were performed.

### Whole cell voltage clamp

#### Extracellular recording solutions

35 mmHg PCO_2_: 124 mM NaCl, 26 mM NaHCO_3_, 1.25 mM NaH_2_PO_4_, 3mM KCl, 1 mM MgSO_4_,10 mM D-glucose, 2 mM CaCl_2_, equilibrated with 5% CO_2_, 95% O_2_, pH 7.4.

55 mmHg PCO_2_: 100 mM NaCl, 50 mM NaHCO_3_, 1.25 mM NaH_2_PO_4_, 3 mM KCl, 1 mM MgSO_4_,10 mM D-glucose, 2 mM CaCl_2_, equilibrated with ∼9% CO_2_ balance being O_2_, pH 7.4.

70 mmHg PCO_2_: 70 mM NaCl, 80 mM NaHCO_3_, 1.25 mM NaH_2_PO_4_, 3 mM KCl, 1 mM MgSO_4_,10 mM D-glucose, 2 mM CaCl_2_, equilibrated with ∼12% CO_2_ balance being O_2_, pH 7.4.

#### Patch recording solution

K-gluconate 130 mM, KCl 10 mM, EGTA 10 mM, CaCl_2_ 2 mM, HEPES 10 mM, sterile filtered, pH adjusted to 7.3 with KOH.

Coverslips containing non-confluent Neuro-2A cells were placed into a perfusion chamber at room temperature standard aCSF. An Axopatch 200B amplifier was used to make whole-cell recordings from the Neuro-2A cells and was controlled by the Strathclyde electrophysiology software programme WinEDR. An agarose salt bridge was used to avoid solution changes altering the potential of the Ag/AgCl reference electrode. All whole-cell recordings were performed at a holding potential of −50 mV. Whole-cell conductance was measured by repeated steps to −30 mV. A typical recording consisted of 2 min at low CO_2_, then 3 min at high CO_2_ and 7 min back to low CO_2_. To allow for any drift in whole-cell conductance unrelated to the CO_2_ stimulus, the maximal conductance during the CO_2_ test stimulus was compared to the average of the conductance before the stimulus and after the stimulus had been fully washed off. The patch clamp voltage clamp recordings were independently analyzed blind to transfection condition.

### Statistical analysis

For the GRAB_ATP_ and whole cell voltage clamp recordings, each cell was considered as an independent replicate. For the cell death assay, each seeding/transfection was considered an independent replicate. Multiple comparisons were analysed with the Kruskal Wallis ANOVA and post hoc Mann Whitney *U*-tests. All quantitative data is presented as box and whisker plots, where the line represents the median, the box is the interquartile range, and the whiskers are the range, with all individual data points included. All statistical tests were performed using R.

## Conflict of Interest

The authors declare that there are no conflicts of interest

## Funding

We thank the Leverhulme Trust (RPG-2022-323, ND) for support. JB was supported by the Biotechnology and Biological Sciences Research Council (BBSRC) and University of Warwick funded Midlands Integrative Biosciences Training Partnership (MIBTP) grant number BB/T00746X/1.

## Data availability

All data generated in the paper is available as supplements to the figures.

---

**Supplementary Figure 1:**
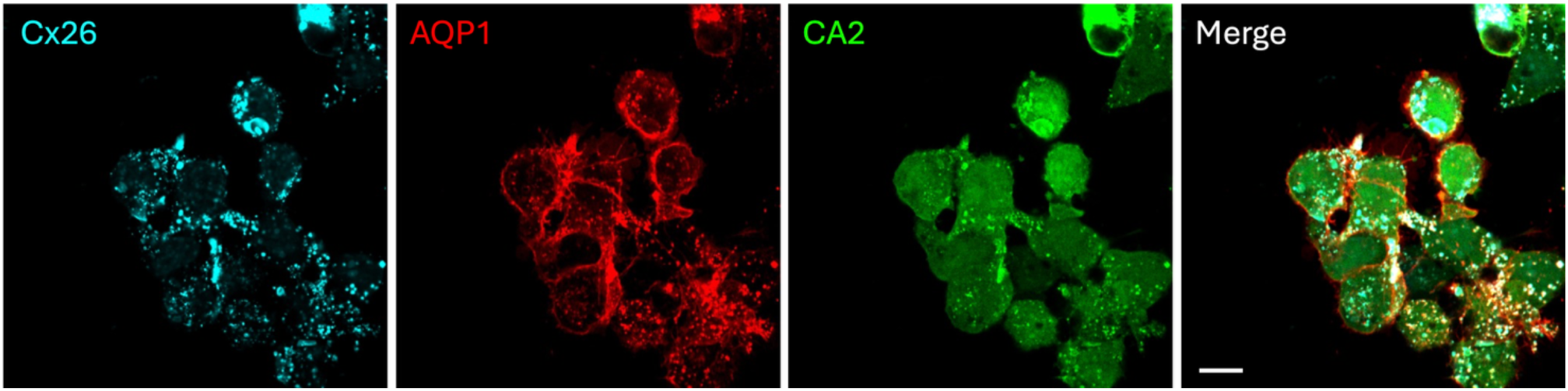
Exemplar images of expression of Cx26, AQP1 and CA2 in N2A cells. Scale bar 10 µm.

**Supplementary Figure 2:**
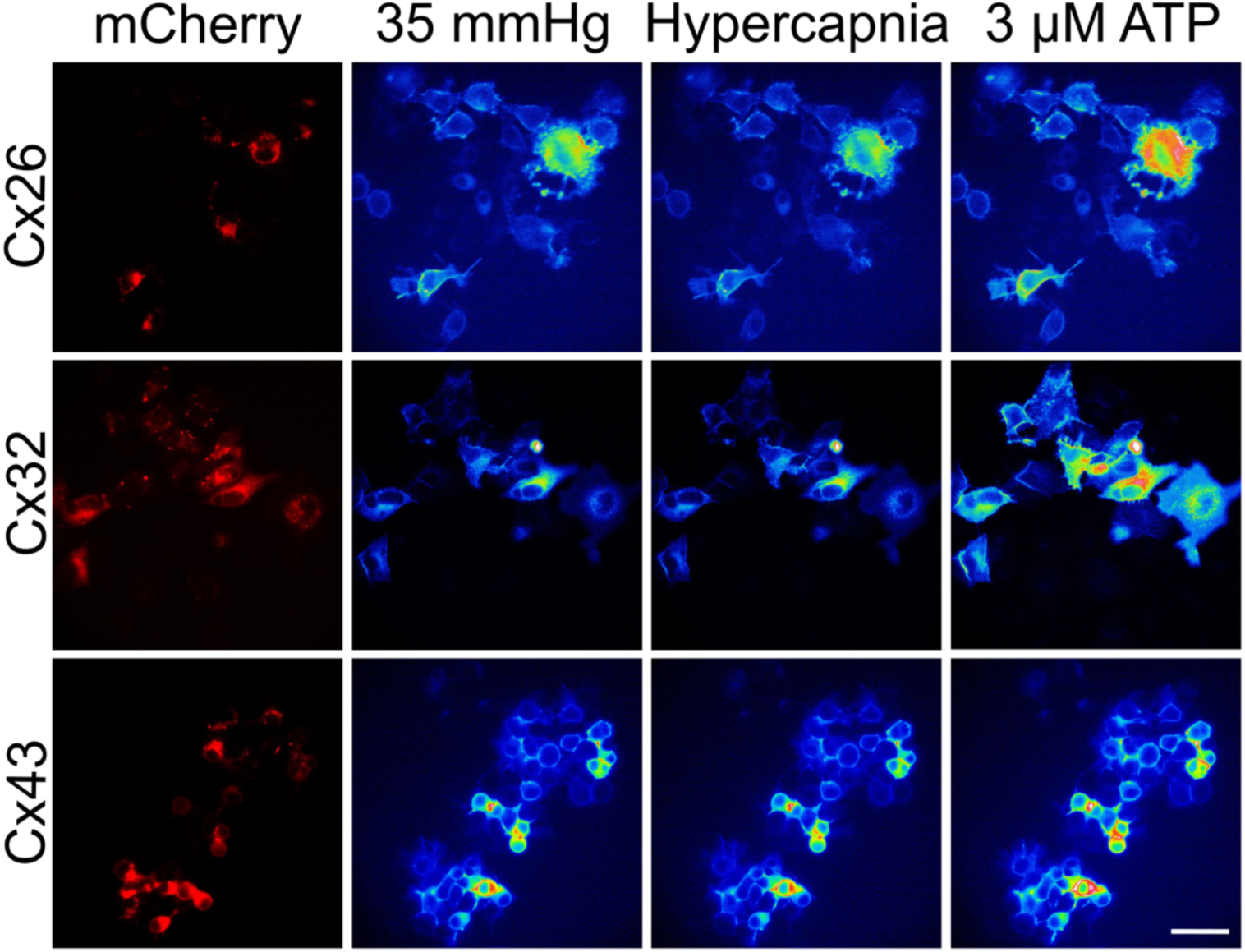
Raw images of connexin expression (mCherry tagged) and GRAB_ATP_ fluorescence for control, hypercapnic conditions (55 mmHg for Cx36 and Cx43, 70 mmHg for Cx32) and 3 µM calibrant. Scale bar 40 µm.

**Supplementary Figure 3:**
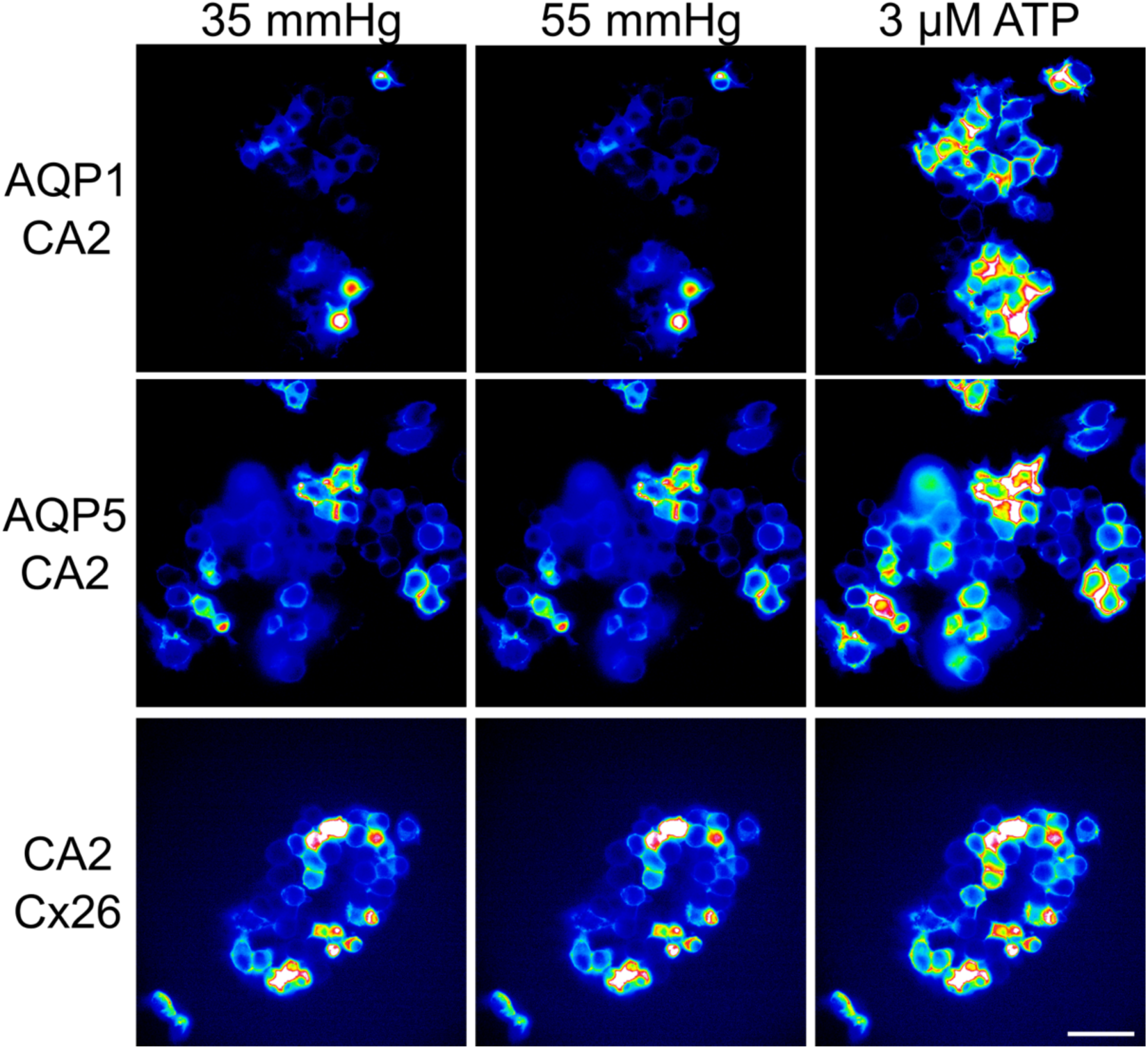
Raw images of GRAB_ATP_ fluorescence from cells transfected with AQP1/CA2, AQP5/CA2 or Cx26/CA2 during control (35 mmHg), hypercapnic conditions (55 mmHg) and 3 µM calibrant. Scale bar 40 µm.

**Supplementary Figure 4:**
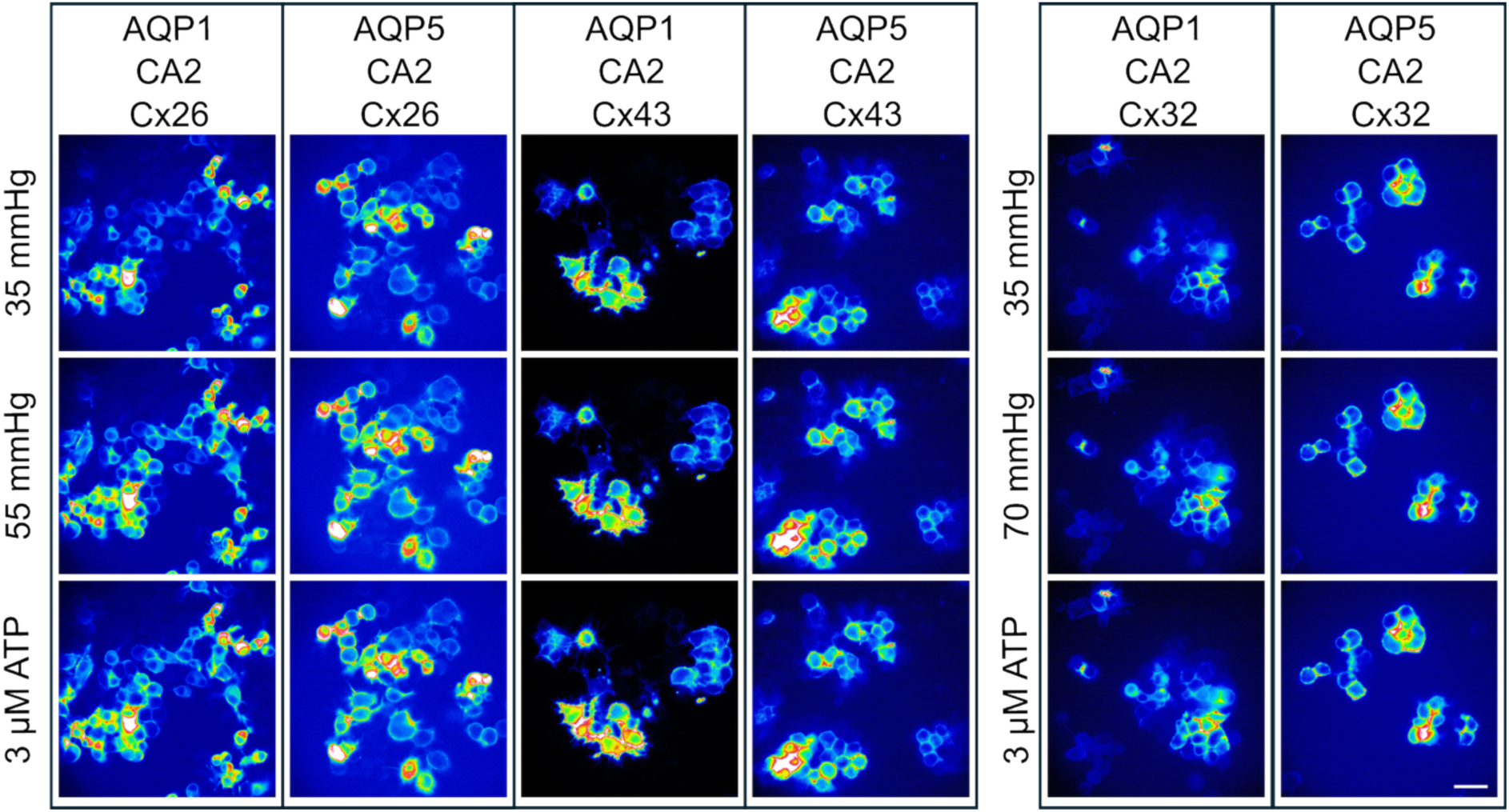
Raw images of GRAB_ATP_ fluorescence from cells transfected with different combinations of CO_2_ sensitive connexins, CO_2_ permeable aquaporins and CA1 during control (35 mmHg), hypercapnic conditions and 3 µM calibrant. Scale bar 40 µm.

## Notes

### Competing Interest Statement

The authors have declared no competing interest.

